# A primitive starfish ancestor from the Early Ordovician of Morocco reveals the origin of crown group Echinodermata

**DOI:** 10.1101/216101

**Authors:** Aaron W. Hunter, Javier Ortega-Hernández

## Abstract

The somasteroids are Ordovician star-shaped animals widely regarded as ancestors of Asterozoa, the group of extant echinoderms that includes brittle stars and starfish. The phylogenetic position of somasteroids makes them critical for understanding the origin and early evolution of crown group Echinodermata. However, the early evolution of asterozoans, the origin of their distinctive body organization and their relationships with other Cambrian and Ordovician echinoderms, such as edrioasteroids, blastozoans, crinoids, and other asterozoans, remain problematic due to the difficulties of comparing the calcitic endoskeleton of these disparate groups. Here we describe the new somasteroid *Cantabrigiaster fezouataensis* from the Early Ordovician (Tremadocian) Fezouata Lagerstätte in Morocco. *Cantabrigiaster* shares with other somasteroids the presence of rod-like virgal ossicles that articulate with the ambulacrals, but differs from all other known asterozoans in the absence of adambulacral ossicles defining the arm margins. The unique arm construction evokes parallels with non-asterozoan echinoderms. Developmentally informed Bayesian and parsimony based phylogenetic analyses, which reflect the homology of the biserial ambulacral ossicles in Paleozoic echinoderms according to the Extraxial-Axial Theory, recover *Cantabrigiaster* as basal within stem group Asterozoa. Our results indicate that *Cantabrigiaster* is the earliest diverging stem group asterozoan, revealing the ancestral morphology of this major clade and clarifying the affinities of problematic Ordovician taxa. Somasteroids are resolved as a paraphyletic grade within stem and crown group Asterozoa (starfishes), whereas stenuroids are paraphyletic within stem group Ophiuroidea (brittle stars). *Cantabrigiaster* also illuminates the relationship between Ordovician crown group Echinodermata and its Cambrian stem lineage, which includes sessile forms with incipient radial symmetry such as edrioasteroids and blastozoans. The contentious Pelmatozoa hypothesis (i.e. monophyly of blastozoans and crinoids) is not supported; instead, blastozoans represent the most likely sister-taxon of crown group Echinodermata.

**Author summary:** Starfish and brittle stars, collectively known as asterozoans, constitute a diverse and ecologically successful group of echinoderms that first appear in the fossil record some 480Ma. However, the early evolution of asterozoans, the origin of their distinctive body organization, and their phylogenetic relationships with Cambrian echinoderms remain largely unresolved. We describe *Cantabrigiaster fezouataensis* gen. et sp. nov., a primitive asterozoan from the Fezouata Lagerstätte, Morocco, with a unique endoskeletal arm organization that reveals the ancestral morphology of this major clade. Bayesian and parsimony based phylogenetic analyses indicate that *Cantabrigiaster* is the earliest diverging stem group asterozoan, and resolve the phylogenetic position of Ordovician asterozoans such as somasteroids. Our analyses clarify the origin of crown group echinoderms relative to their problematic Cambrian stem group representatives.

## Introduction

Asterozoans – whose most familiar members include starfish and brittle stars – are the dominant group of extant echinoderms based on their diversity, abundance, and biogeographic distribution [1]. Despite their ecological success and a fossil record spanning more than 480 million years [2–4], the origin and early evolution of asterozoans, and that of crown group echinoderms more generally, remains uncertain given the difficulty of comparing the organization of the calcified endoskeleton in diverse groups of Lower Paleozoic ancestors, such as the edrioasteroids and blastozoans [5–13]. The Extraxial-Axial Theory (EAT), which supports the homology of the biserial ambulacral ossicles of pentarradial echinoderms based on embryonic and ontogenetic data [14–16], has been proposed as a developmentally-informed model that would facilitate making comparisons among groups with disparate morphologies. Although the EAT can potentially clarify the early evolution of crown group Echinodermata, the broad implications of this hypothesis have never been examined under a comprehensive phylogenetic framework. Consequently, the main phylogenetic predictions of the EAT pertaining to the evolutionary relationships of Cambrian and Ordovician echinoderms, such as the origin of the crown group from edrioasteroid-like ancestors [14–16, 17], have yet to be critically tested.

Here, we describe the new somasteroid *Cantabrigiaster fezouataensis* gen. et sp. nov. from the Early Ordovician (Tremadocian) Fezouata Shale in Zagora, central Anti-Atlas, Morocco [4] (Fig S1 and SI text). The exceptionally preserved morphology of *Cantabrigiaster* reveals a unique organization among somasteroids, and allows us to test the phylogenetic implications of this taxon for the origin of Asterozoa and crown group Echinodermata.

## Results

### Systematic Paleontology

(crown group) Echinodermata Bruguière, 1791

(stem group) Asterozoa Zittel, 1895

Somasteroidea Spencer, 1951

*Cantabrigiaster fezouataensis* gen. et sp. nov.

#### Etymology

Genus name derived from ‘*Cantabrigia’*, after the cities of Cambridge in the UK and USA, which were home to the influential asterozoan workers John William Salter (University of Cambridge), Juliet Shackleton (neé Dean) (University of Cambridge), and Howard Barraclough ‘Barry’ Fell (Harvard University).

#### Holotype

FSL, VOMN 424 961 (Fig 1).

**Fig 1.**
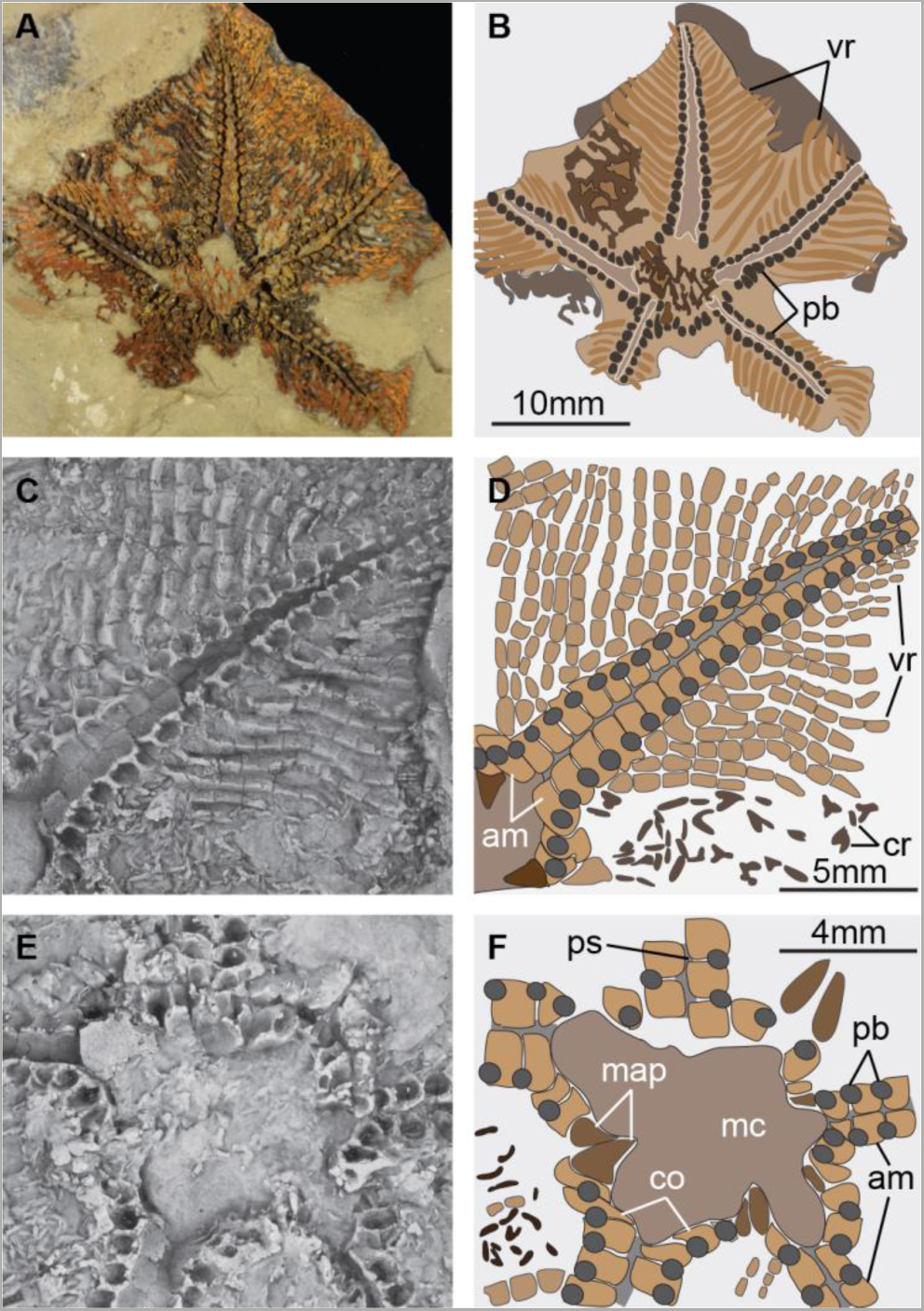
*Cantabrigiaster fezouataensis* from the Lower Ordovician (Tremadocian) of Morocco. Holotype FSL-VOMN-424961. (A) Oral view (body fossil). (B) Interpretative diagram of A. (C) Close-up of extended arm (latex mould). (D) Interpretative diagram of C. (E) Close-up of oral region (latex mould). (F) Interpretative diagram of E. Abbreviations: am, ambulacral ossicles; co, circumoral ossicles; cr, carinal region ossicles; map, mouth angle plates; mc, mouth cavity; pb, podial basins; ps, podial suture; vr, virgal ossicles.

#### Referred material

YPM IP 535545-535559 (Fig S2 and Fig S3).

#### Diagnosis for genus and species

Somasteroid typified by biserial and offset ambulacrals with thin transverse bar, wide perradial groove, multiple interconnected virgal ossicles, and aboral carinal region with network of spicule-like ossicles. Adambulacral ossicle series lacking along abaxial body margins.

#### Description

The arms are broad, petaloid, and arranged in a pentagonal outline (Figs 1A, 1B, S3A, S3C and S3D). The aboral skeleton (carinal region) is composed of randomly scattered spicule-like ossicles arranged into an irregular network (Figs 1A, 1B, S2D, S3E and S3G). On the oral side, the ambulacrals consist of flattened ossicles with a subquadrate outline. These ossicles abut each other following the orientation of the perradial axis (Figs 1C, 1D, S2A, S2C, S2F and S2G). The perradial suture is straight, and the ambulacrals at either side are stepped out of phase by approximately half an ossicle. The abaxial organization of the ambulacrals consists of an elevated perradial ridge, less than a quarter in width relative to the ambulacral, and bears a thin transverse bar that occupies a central position conferring a T-shape in oral view (Figs 1C, 1D, S2A, S2E and S2G). The perradial ridges of the ambulacral ossicles at either side of the perradial suture are substantially separated from each other, forming a wide oral groove (Figs 1C-E, S2A-C and S2G). The podial basins are shared equally between adjacent ambulacrals. Abaxially, the following ossicle series consist of the perpendiculars, also known as virgals in somasteroids [2, 5, 6]. The perpendicular series is composed of interconnected and robust rod-like virgal ossicles without spines. These ossicles follow a perpendicular orientation relative to the perradial suture (Figs 1A-D and S2). The virgal ossicles close to the ambulacrals are the largest, and become smaller in length and width towards the abaxial body margins. Likewise, adjacent perpendicular series are in direct contact with each other adaxially relative to the perradial suture, whereas it is possible to observe open gaps between them towards the abaxial body margins. Proximal (relative to the mouth) perpendiculars series consist of up to nine virgal ossicles, that gradually decrease in number towards the tips of the arms (Figs 1A-D and 2). The circumoral ossicles are enlarged relative to ambulacral ossicles, and the first podial pore is shared equally with the small and sub-triangular mouth angle plates (Figs 1E and 1F). The madreporite is not preserved.

## Discussion

The presence of virgal ossicles in *Cantabrigiaster* strongly supports its affinities with somasteroids [2, 5–9, 14, 21]. *Cantabrigiaster* bears the greatest similarity to the Tremadocian taxa *Chinianaster*, *Thoralaster*, and *Villebrunaster* (Fig S4), but is unique among somasteroids in lacking ossicles along the abaxial lateral margins of the arms (Figs 1A and 1D). The arm construction of *Cantabrigiaster* consists of flattened and offset biserial ambulacrals, each of which articulates with an abaxially-oriented (i.e. perpendicular) perpendiculars series composed of simple virgal ossicles (Fig 2). In addition to these features, the arms of all other somasteroids also possess a series of axially-oriented ossicles along the lateral margins that vary from small and bead-like – albeit with occasional spikes – in Tremadocian taxa [2, 5, 6] (Fig S4), to robust and block-like in the stratigraphically younger (Floian) *Ophioxenikos* [10] and *Archegonaster* [9]. These comparisons suggest a selective pressure towards the addition of new ossicle series among early asterozoans (Fig S5). *Cantabrigiaster* embodies the ancestral condition by virtue of lacking ossicles defining the lateral arm margins (Figs 1 and 2), whereas other somasteroids record the first appearance of these structures along the edges of the arms, and their subsequent changes in size and shape. Based on this sequence, we propose that the origin of new axially-oriented ossicle series in early asterozoans required their formation on the abaxial edges of the arms. Our hypothesis implies that the proximity of axially-oriented ossicle series relative to the perradial axis reflects the order of their evolutionary appearance (Fig S5 and SI text); since virgals are abaxially-oriented, they are not directly comparable with any of the axially-oriented ossicle series observed in Paleozoic asterozoans. In this context, *Cantabrigiaster* specifically lacks the adambulacral ossicle series present in more derived somasteroids, stenuroids, ophiuroids and asteroids, highlighting its profound significance for understanding the evolution of the asterozoan body plan.

**Fig 2.**
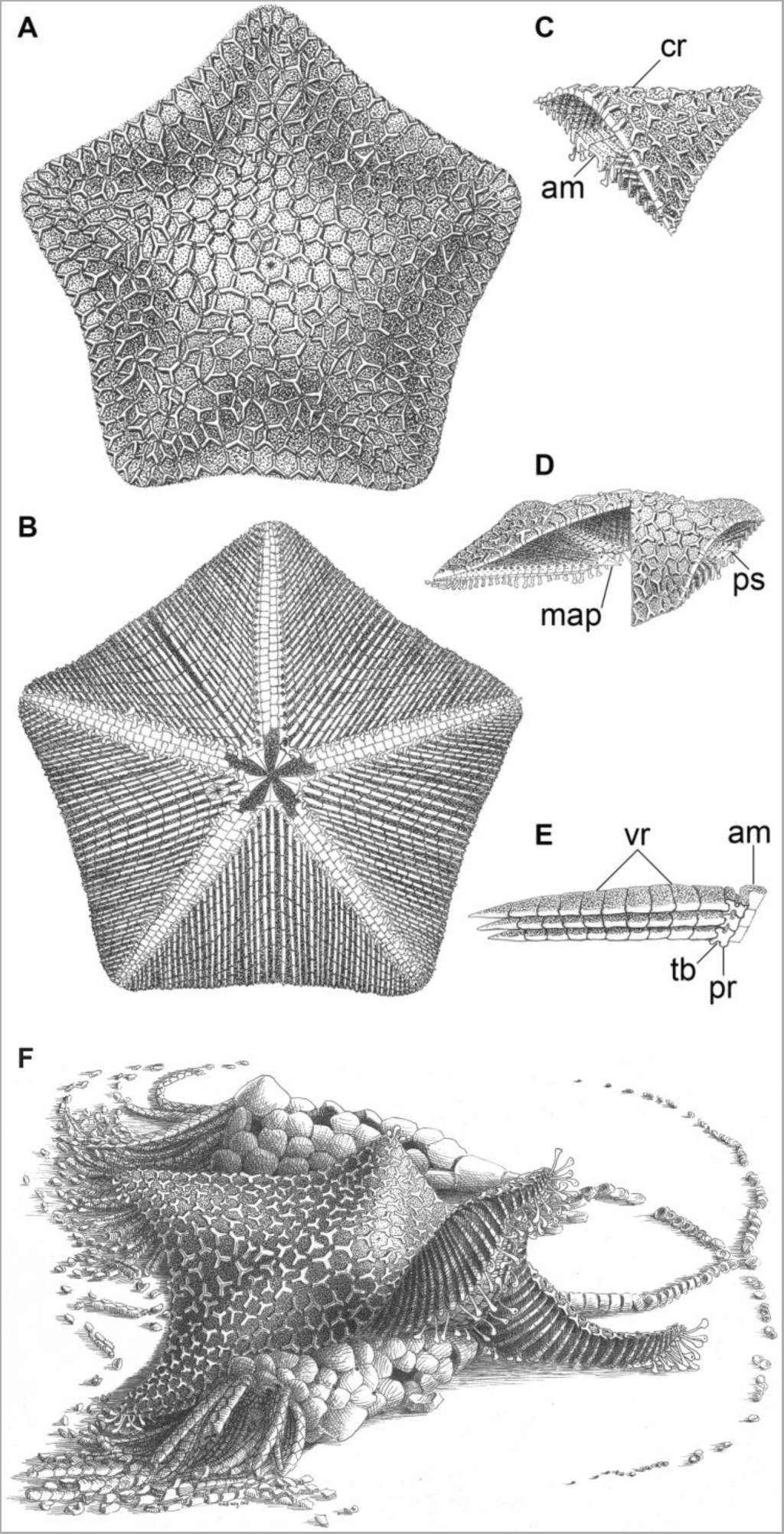
Morphological reconstruction of *Cantabrigiaster fezouataensis.* (A) Aboral view. (B) Oral view. (C) Cross section of isolated arm in oblique view. (D) Cross section of main body cavity lateral view. (E) Isolated virgal ossicle series and ambulacrals in oral view. (F) Life reconstruction of *Cantabrigiaster fezouataensis*. Artwork by Marguerite Lardanchet. Abbreviations: am, ambulacral ossicles; cr, carinal region ossicles; map, mouth angle plates; pr, perradial ridge; ps, podial suture; tb, transverse bar; vr, virgal ossicles.

The Extraxial-Axial Theory (EAT) supports the homology of the ambulacrals across pentaradial total-group echinoderms based on their developmental origin and postembryonic ontogeny [14–17], and allows comparison of the skeletal organization of *Cantabrigiaster* in a broader phylogenetic scale. Outside Asterozoa, the absence of adambulacrals in *Cantabrigiaster* draws parallels with Tremadocian crinoids (e.g. protocrinoids, *Apektocrinus, Eknomocrinus*), whose arm construction incorporates flattened and offset biserial ambulacrals articulated to an abaxially-oriented series of simple ossicles, here expressed as the cover plates [13, 16–18] (Fig S5 and SI text). A similar axial skeletal organization is also observed among Cambrian forms, most notably edrioasteroids – which also possess flattened and offset biserial ambulacrals but lack free appendages [9, 11, 19], and to a lesser extent blastozoans, which have free appendages formed by modified ambulacrals known as brachioles [12, 20, 21]. The widespread occurrence of these characters among non-asterozoan groups suggests that their presence in *Cantabrigiaster* is symplesiomorphic.

We designed a comprehensive phylogenetic analysis of Lower Paleozoic total-group echinoderms in order to test the significance of *Cantabrigiaster* for the origin of Asterozoa. The dataset reflects the ambulacral homology proposed by the EAT [14–18], the oral symmetry model proposed by Universal Element Homology [22–24], and our hypothesis for the correspondence of axially-oriented ossicle series in early asterozoans (Fig S5 and SI text). Bayesian and parsimony-based analyses recover practically identical topologies (Figs 3, S6 and S7), despite a modest loss in tree resolution that can be expected from the former methodology, indicating a robust phylogenetic signal [25]. *Cantabrigiaster* occupies a basal position within total-group Asterozoa, supporting our hypothesis that the absence of adambulacrals is ancestral. Tremadocian somasteroids are resolved as a pharaphyletic grade of stem group asterozoans (per refs [2, 26]; *contra* ref. [5]), whereas the Floian *Ophioxenikos* [10] and *Archegonaster* [9] consistently occupy a more derived position as members of crown group Asterozoa. The analyses argue against the monophyly of stenuroids [6], but corroborate their close phylogenetic relationship to ophiuroids, specifically as their earliest diverging stem group representatives [2, 5, 7, 26]. These findings indicate that the evolution of a well-developed adambulacral ossicle series constitutes a critical step in the origin of crown group Asterozoa, and demonstrate that the abaxially-oriented virgals of somasteroids became independently reduced – and ultimately lost – within the stem lineages of Ophiuroidea and Asteroidea [6] (Fig S5).

**Fig 3.**
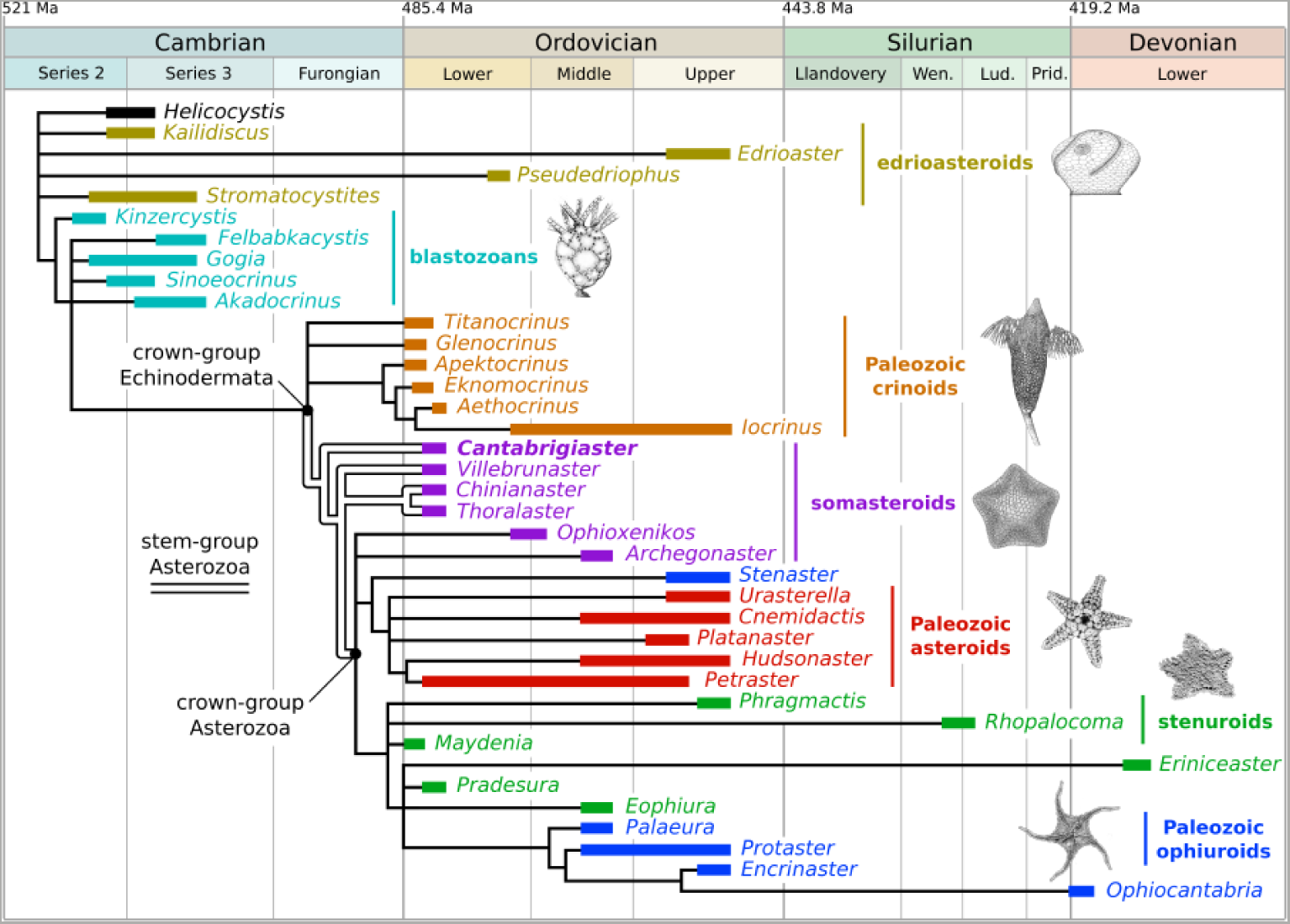
Evolution of crown group Echinodermata. Consensus topology based on the Bayesian-inference analysis of 38 taxa and 73 morphological characters informed by the EAT [14, 15] (SI text). See Fig S6 for support values and comparison with the results of the parsimony-based phylogenetic analyses. Stratigraphic ranges of taxa based on refs [3, 6, 12, 18, 27].

Our results also clarify the heated debate over the phylogenetic placement of Cambrian edrioasteroids and blastozoans relative to Ordovician crinoids and asterozoans [9, 11, 13–18, 20, 22–24, 27] (Figs 3, S6 and S7). Edrioasteroids and blastozoans are resolved as stem group echinoderms. Contrary to previous hypotheses [8, 9, 11] and predictions from EAT proponents [13, 16–18], blastozoans – rather than edrioasteroids – are the most derived stem group representatives, making them strong candidates for the sister-taxon of crown group Echinodermata. This position confirms that blastozoans are ancestral relative to crinoids, and simultaneously falsifies the monophyly of these taxa according to the Pelmatozoa hypothesis [11, 20, 22–24, 27]. Character mapping indicates that most of the features that Tremadocian crinoids share with edrioasteroids (e.g. flattened and offset biserial ambulacrals, cover plates [15–18]) and blastozoans (e.g. irregular thecal plating, extended perforate region, 2-1-2 symmetry [22, 23]) are symplesiomorphic (Fig S5). The consensus topology suggests a single origin for the free appendages of blastozoans and crown group echinoderms, albeit with fundamental differences in their endoskeletal construction [14,16,17]. Brachioles are exclusive – and most likely autapomorphic – to blastozoans [12, 20, 21]. The presence of a straight perradial suture, and the aboral extension of the body wall over the arms forming coelomic cavities, represent fundamental synapomorphies uniting total-group Crinoidea and total-group Asterozoa [15–17], despite rare examples of convergence within the echinoderm stem group [22, 27]. Ultimately, our findings reconcile the evidence supporting the homology of ambulacral and oral ossicle organization in edrioasteroids, blastozoans, crinoids and asterozoans, into a robust phylogenetic hypothesis that informs the origin of crown group Echinodermata and the gradual early evolution of the archetypical asterozoan body plan (Figs 3 and S5).

## Materials and Methods

### Specimen analysis

Studied material is deposited at the Faculty of Science, Claude Bernard University of Lyon 1 (FSL-VOMN), Natural History Museum of Nantes (MHNN), National Museum, Prague (NM-P), and Yale Peabody Museum, Yale University (YPM). Latex molds were made of all the material with the exception of that of the YPM. The material was photographed with a Nikon D5500 SLR fitted with Micro Nikkor 40mm.

### Phylogenetic analysis

The character matrix for the phylogenetic analyses includes 38 taxa and 73 characters (see Dataset S1 and S2); detailed discussion of character scoring and applicability are provided below. The Bayesian analysis was run in MrBayes 3.2 using the Monte Carlo Markov-chain model for discrete morphological characters [28, 29] for 10 million generations (four chains), with every 1000^th^ sample stored (resulting in 10,000 samples), and 25% burn-in (resulting in 7,500 retained samples). The parsimony analyses were run in TNT [30] under New Technology Search, using Driven Search with Sectorial Search, Ratchet, Drift, and Tree fusing options activated with standard settings [31, 32]. The analysis was set to find the minimum tree length 100 times and to collapse trees after each search. All characters were treated as unordered. For comparative purposes, analyses were performed under equal and implied weights (*k*=3) to test the effect of homoplasy penalization on the position of *Cantabrigiaster* and the robustness of the dataset [33]. Comparisons between results of the phylogenetic analyses are presented in figures S6 and S7. Parsimony-based analysis under Traditional Search with 10,000 replicates produced identical results as those obtained under New Technology Search.

## Acknowledgements

We acknowledge support from a Herchel Smith Research Fellowship in Biological Sciences, a Bye-Fellowship at Emmanuel College (both JO-H) and visiting fellowship at Clare Hall (AWH). National Geographic for funded the collection of the holotype. Dr. Emmanuel Robert (FSL) is thanked for assisting access to the holotype and other figured material. Additional thanks go to Dr. Martin Valent (NM-P) for access to types of *Archegonaster*. Dr. Fred Hotchkiss (MPRI) and the Yale Peabody Museum assisted in securing the lectotypes.

## Notes

The authors declare no competing interests.

